# Precise annotation of tick mitochondrial genomes reveals multiple STR variation and one transposon-like element

**DOI:** 10.1101/753541

**Authors:** Ze Chen, Xiaofeng Xu, Xiaolong Yang, Zhijun Yu, Yonghong Hu, Duo Wang, Wenjun Bu, Jingze Liu, Shan Gao

## Abstract

In this study, we used long-PCR amplification combined with Next-Generation Sequencing (NGS) to obtain complete mitochondrial genomes of individual ticks and performed precise annotation of these genomes. These annotations were confirmed by the PacBio full-length transcriptome data to cover both entire strands of the mitochondrial genomes without any gaps or overlaps. Based on these annotations, most of our findings were consistent with those from previous studies. Moreover, two important findings were reported for the first time, contributing fundamental knowledge to mitochondrial biology. The first was the discovery of a transposon-like element that may reveal the mechanisms of mitochondrial gene order rearrangement and genomic structural variation. Another finding was that Short Tandem Repeat (STRs) are the dominant variation type causing mitochondrial sequence diversity within an individual tick, insect, mouse and human. Comparisons between interindividual and intraindividual variation showed that polynucleotides and STRs with longer repeat units had the same variation pattern. Particularly, mitochondria containing deleterious mutations can accumulate in cells and deleterious STR mutations irreversibly change the proteins made from their mRNAs. Therefore, we proposed that deleterious STR mutations in mitochondria cause aging and diseases. This finding helped to ultimately reveal the mechanisms of mitochondrial DNA variation and its consequences (e.g., aging and diseases) in animals. Our study will give rise to the reconsideration of the importance of STRs and a unified study of STR variation with longer and shorter repeat units (particularly polynucleotides) in both nuclear and mitochondrial genomes. The complete mitochondrial genome sequence of *Dermacentor silvarum* is available at the NCBI GenBank database under the accession number MN347015 and the raw data is available at the NCBI SRA database under the accession number SRP178347.

## Introduction

Mitochondrial genome annotation is indispensable for fundamental research in numerous fields, including mitochondrial biochemistry, physiology, and the molecular phylogenetics and evolution of animals. Moreover, high-resolution annotation of animal mitochondrial genomes can be used to investigate RNA processing, maturation, degradation, and even the regulation of gene expression [1]. Two substantial contributions to the improvement of mitochondrial genome annotation have been published. Gao *et al*. constructed the first quantitative transcription map of animal mitochondrial genomes by sequencing the full-length transcriptome of the insect *Erthesina fullo* Thunberg [1] on the PacBio platform [2]. New findings included the 3′ polyadenylation and possible 5′ m^7^G caps of rRNAs [1], the polycistronic transcripts [1], the antisense transcripts of all mitochondrial genes [1], and novel long non-coding RNAs (lncRNAs) [3]. Based on these findings, we proposed the mitochondrial cleavage model [3] and the mitochondrial tRNA processing model [4]. In addition, we proposed that although all antisense transcripts are processed from two primary transcripts, long antisense transcripts degrade quickly as transient RNAs, making them unlikely to perform specific functions [4]. The second contribution involved the use 5′ and 3′ end small RNAs (5′ and 3′ sRNAs) [4] to annotate mitochondrial genes at 1 bp resolution, subsequently dubbed precise annotation [5]. Using these accurate genomes with precise annotations, we discovered that a novel 31-nt ncRNA exists in mammalian mitochondrial DNA [4] and that the copy numbers of tandem repeats exhibit great diversity within an *E. fullo* individual [5]. Recently, precise annotation of human, chimpanzee, rhesus macaque and mouse mitochondrial genomes has been performed to investigate five Conserved Sequence Blocks (CSBs) in the mitochondrial D-loop region [6], which ultimately led to an understanding of the mechanisms involved in the RNA-DNA transition and even the functions of the D-loop.

In this study, we used long-PCR amplification combined with Next-Generation Sequencing (NGS) to obtain complete mitochondrial genomes of individual ticks and performed precise annotation of these genomes. Given that mtDNA isolation and purification are circumvented in our method and in the Whole-Genome Sequencing (WGS) method, both are simple and cost-effective. However, compared to the WGS method, our method has a number of advantages: (i) errors in the assembly of mitochondrial genomes caused by highly similar exogenous or nuclear sequences [i.e., Nuclear Mitochondrial DNA (NUMT)] are avoided; (ii) highly similar segments (e.g., control regions 1 and 2 of *D. silvarum*) of mitochondrial genomes can be assembled separately (**Results**); (iii) using high-depth sequencing, the sequence heterogeneity and DNA variation in mitochondrial genomes within an individual can be accurately determined. In this study, we aimed to achieve the following research goals: (1) to provide a simple, cost-effective and accurate method for the study of extremely high AT-content mitochondrial genomes within an individual animal containing miniscule DNA. (2) to provide a high-quality reference genome with precise annotation for future studies of tick mitochondrial genomes; and (3) to use these mitochondrial genomes containing two Control Regions (CRs) to confirm our previous findings [5] from those containing one CR.

## Results

### Using Long-PCR and NGS to obtain complete mitochondrial genomes of individual ticks

A previous study classified tick mitochondrial genomes into three types: I, II and III for Argasidae (soft tick), Prostriata (Ixodes spp.), and Metastriata (all other hard ticks), respectively, based on their gene orders [7] (**Figure 1A**). The present study focused on the genus Dermacentor belonging to Metastriata using ticks from five species (*D. silvarum, D. nuttalli, D. marginatus, D. niveus*, and *D. ushakovae*). The type III mitochondrial genomes of individual ticks (**Figure 1B**) were obtained using long-PCR amplification combined with NGS (**Materials and Methods**). All the reference genomes of tick mitochondria read in the 5’→3’ direction as the major coding strand (J-strand). Using specific primers (**Table 1**), each complete mitochondrial genome was amplified in two large segments: large segment 1 (L1) and large segment 2 (L2) or large segment 3 (L3) and large segment 4 (L4). L1 and L2 contain Control Region 1 (CR1) and Control Region 2 (CR2), respectively, while L3 and L4 contain tandem Repeat 1 (R1) and tandem Repeat 2 (R2), respectively (**Figure 1A**). Using ∼4 Gbp 2 × 150 DNA-seq data for each genome, the complete mitochondrial genomes of *D. silvarum, D. nuttalli* and *D. marginatus* were obtained by assembling L3 and L4 separately then merging L3 and L4 (**Figure 1B**). In addition, CR1 and CR2 on L4 were validated using PCR amplification combined with Sanger sequencing, separately, as CR1 and CR2 share an identical segment (**Figure 2A**). Using ∼4 Gbp 2 × 150 DNA-seq data, the complete mitochondrial genome of *D. silvarum* was also obtained by assembling L1 and L2 separately then merging L1 and L2. Furthermore, R1 and R2 on L1 were validated using PCR amplification combined with Sanger sequencing, separately, as the repeat units of R1 are the reverse complements of the repeat units of R2 (**Figure 3**). A comparison was performed between the *D. silvarum* mitochondrial genome obtained by sequencing L3 and L4 and that by sequencing L1 and L2 to improve the accuracy of its DNA sequence. As both R1 and R2 are longer than 150 bp, we also used ∼4 Gbp 2 × 250 bp DNA-seq data to obtain full-length sequences of R1 and R2 directly for genome polishing. In total, 12.7 Gbp DNA-seq data were generated to cover ∼848,069 × (12.72 Gbp/1.5 Kbp) of the *D. silvarum* mitochondrial genome (GenBank: MN347015) which was ultimately used as a reference for precise annotation in the following studies.

**Table 1.**
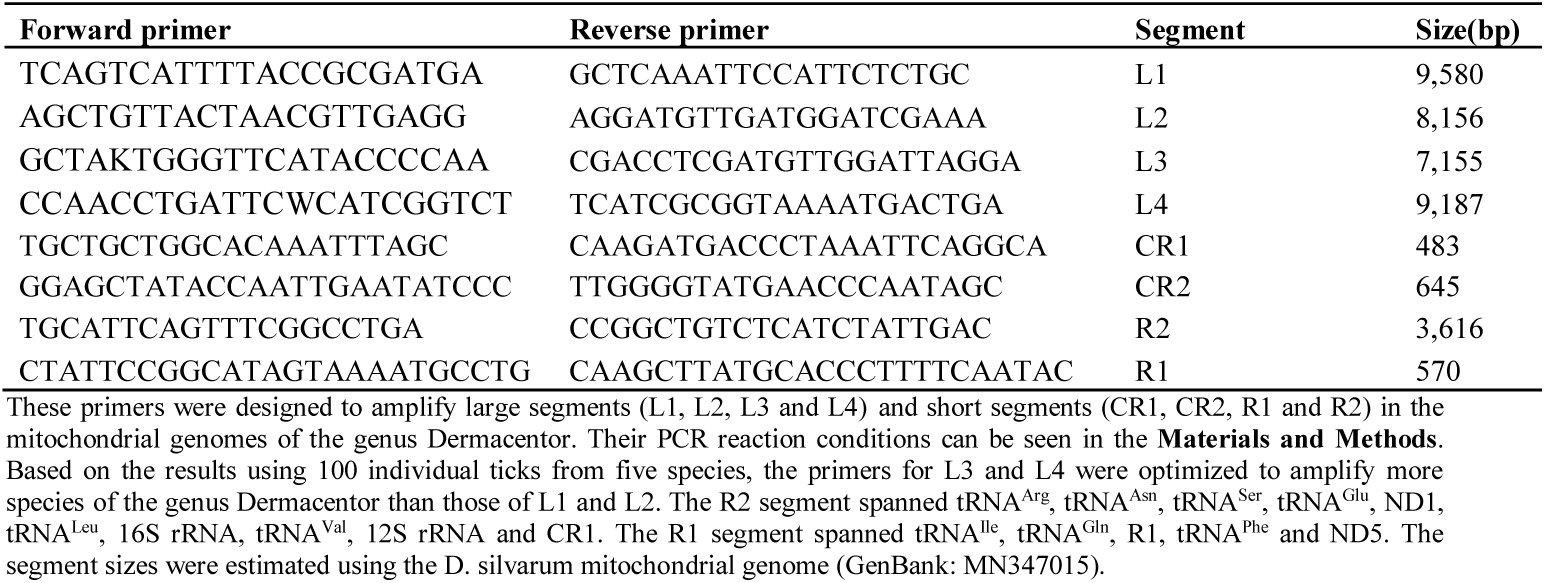
PCR primers for the *Dermacentor* mitochondrial genomes

**Figure 1.**
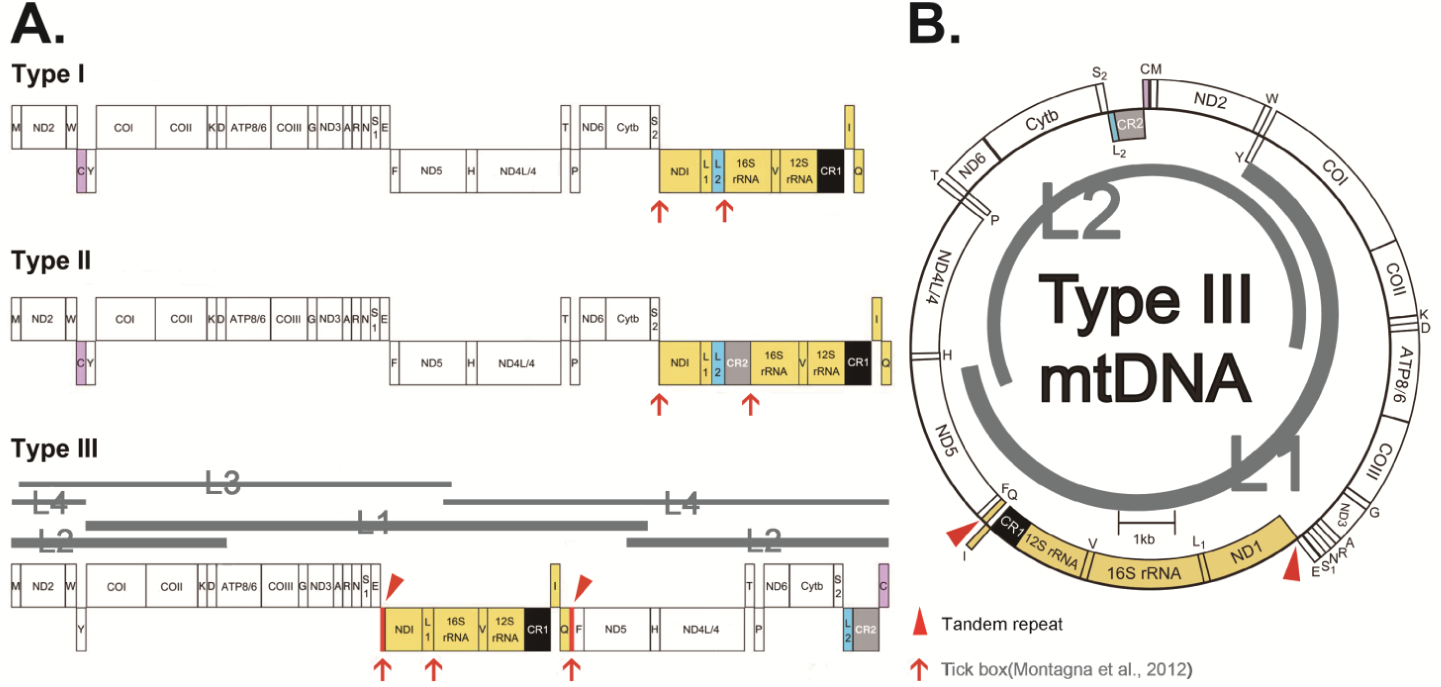
Long PCR amplification of each entire mitochondrial genome. All the primers and PCR reaction conditions are listed in Table 1. The tRNA genes are represented by their single letter codes. CR1 and CR2 represents the control region 1 and the control region 2, respectively. Translocated genes are reported in the same colour. All the reference genomes of tick mitochondria read in the 5’→3’ direction as the major coding strand (J-strand). **A**. The genes from the J-strand and the N-strand are deployed upward and downward, respectively. The tick mitochondrial genomes were classified into three types, which are type I, II and III (Results). The type III mitochondrial genomes can be amplified into two large segments (L1&L2 or L3&L4) by long-PCR using total DNA from individual ticks. Using the complete *D. silvarum* mitochondrial genome (GenBank: MN347015), L1, L2, L3 and L4 were estimated as ∼9.6, ∼8.2, ∼7.2 and ∼9.2 Kbp in size (**Table 1**), respectively. **B**. The gene order of the type III mitochondrial genomes of ticks. The genes from the J-strand and N-strand are deployed outward and inward, respectively.

**Figure 2.**
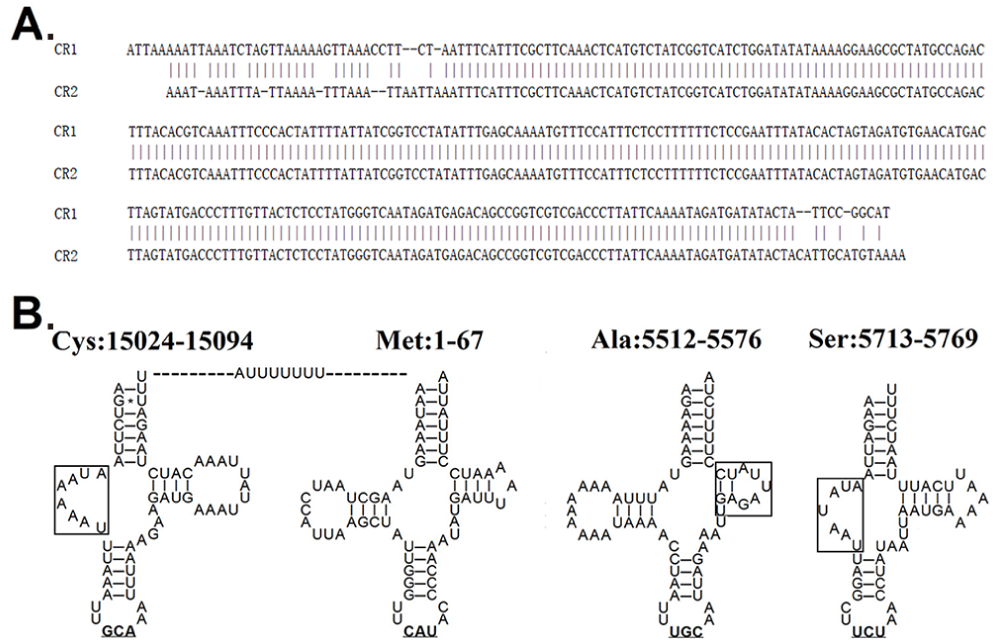
Precise annotation of mitochondrial tRNAs and control regions. A. CR1 and CR2 were determined in the D. silvarum mitochondrial genome (GenBank: MN347015). B. In MN347015, small RNA A[U]7 was produced from between tRNACys and tRNAMet. One of tRNASer and tRNACys had no D-arm, while tRNAAla, tRNAGlu, tRNATyr and tRNAPhe had unstable T-arms (indicated in black box).

**Figure 3.**
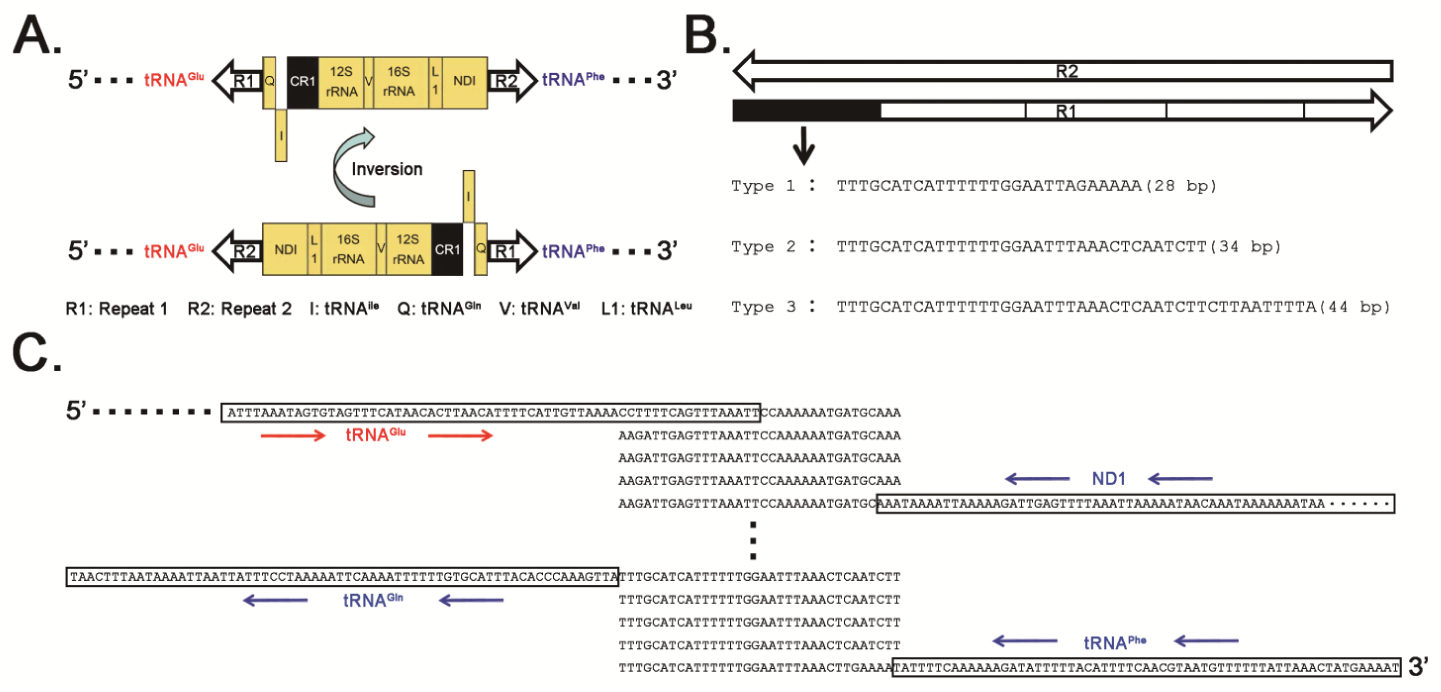
The transposon-like element in the D. silvarum mitochondrial genome. All the mitochondrial genomes read in the 5’→3’ direction as the J-strand. The genes from the J-strand and the N-strand are indicated in red and blue colours, respectively. A. The genes from the J-strand and the N-strand are deployed upward and downward, respectively. B. R1 and R2 were composed of several repeat units, respectively. And the repeat units in R1 are reverse complimentary to those in R2. In total, three types of repeat units (type 1, 2 and 3) of R1 were identified. C. R1 and R2 were determined to have 5 repeat units in the D. silvarum mitochondrial genome (GenBank: MN347015).

Comparison of *D. silvarum, D. nuttalli* and *D. marginatus* mitochondrial genomes showed that they have the same gene order (type III) and high sequence identities (> 95%) without considering differences in the copy numbers of tandem repeats. Preliminary analysis showed two significant features in these tick mitochondrial genomes that are also possible in ticks of Metastriata (**Figure 1A**): 1) these tick mitochondrial genomes contained two tandem repeats (R1 and R2) and 2) they contained multiple Short Tandem Repeats (STRs) with very short repeat units (1 or 2 bp). STRs, widely used by forensic geneticists and in genetic genealogy, are often referred to as Simple Sequence Repeats (SSRs) by plant geneticists [4] or microsatellites. Found widely in animal mitochondrial genomes, STRs follow a pattern in which one or more nucleotides (repeat unit) are repeated and the repeat units are directly adjacent to each other allowing for very rare Single Nucleotide Polymorphisms (SNPs) in the repeat units. The minimum length of the repeat units in STRs is 1 bp; this type of STR is a polynucleotide. PolyAs and polyTs occur frequently in the tick and in insect mitochondrial genomes, contributing substantially to the high AT content. Another reason for a unified study STR variation with longer and shorter repeat units (particularly polynucleotides) was that polynucleotides exhibited the same variation pattern as tandem repeats R1 and R2 in the *D. silvarum* mitochondrial genome (see the paragraphs to follow). To briefly describe a tandem repeat, we use the repeat unit and its copy number. STRs can be classified by their repeat unit length (m) and copy number (n), thus noted as m×n STR. For example, the STR ATATATATAT is noted as [AT]_5_ and classified as 2×5 STR. In this way, a polynucleotide is classified as 1×n STR.

### Precise annotation of the *D. silvarum* mitochondrial genome

We performed precise annotation of the complete *D. silvarum* mitochondrial genome (**Table 2**) using sRNA-seq data and confirmed these annotations using the PacBio full-length transcriptome data (**Materials and Methods**). Although this tick’s mitochondrial genome contains two CRs (CR1 and CR2), unlike those in most other animals, the precise annotation of the *D. silvarum* mitochondrial genome still confirmed our previous findings, particularly the mitochondrial cleavage model [3] and the mitochondrial tRNA processing model [4]. *D. silvarum* transcribes both entire strands of its mitochondrial genome to produce two primary transcripts covering CR1 and CR2, predicted to be non-coding and non-transcriptional regions in the previous study [7]. CR1 with a length of 309 bp and CR2 with a length of 307 bp shared an 263-bp identical segment (**Figure 2A**). CR1 and R1 were annotated as full-length RNAs cleaved from the minor coding strand (N-strand) primary transcript, while CR2 and R2 were annotated as DNA regions (**Table 2**) covered by four transient RNAs. Our *D. silvarum* mitochondrial genome shared a sequence identity of 97.47% with the publicly available *D. silvarum* mitochondrial genome NC_026552.1 in the NCBI RefSeq database. Although most of the new annotations were consistent with those of NC_026552.1, we corrected many errors in NC_026552.1, particularly in tRNAs, rRNAs, CR1, CR2, R1 and R2.

**Table 2.**
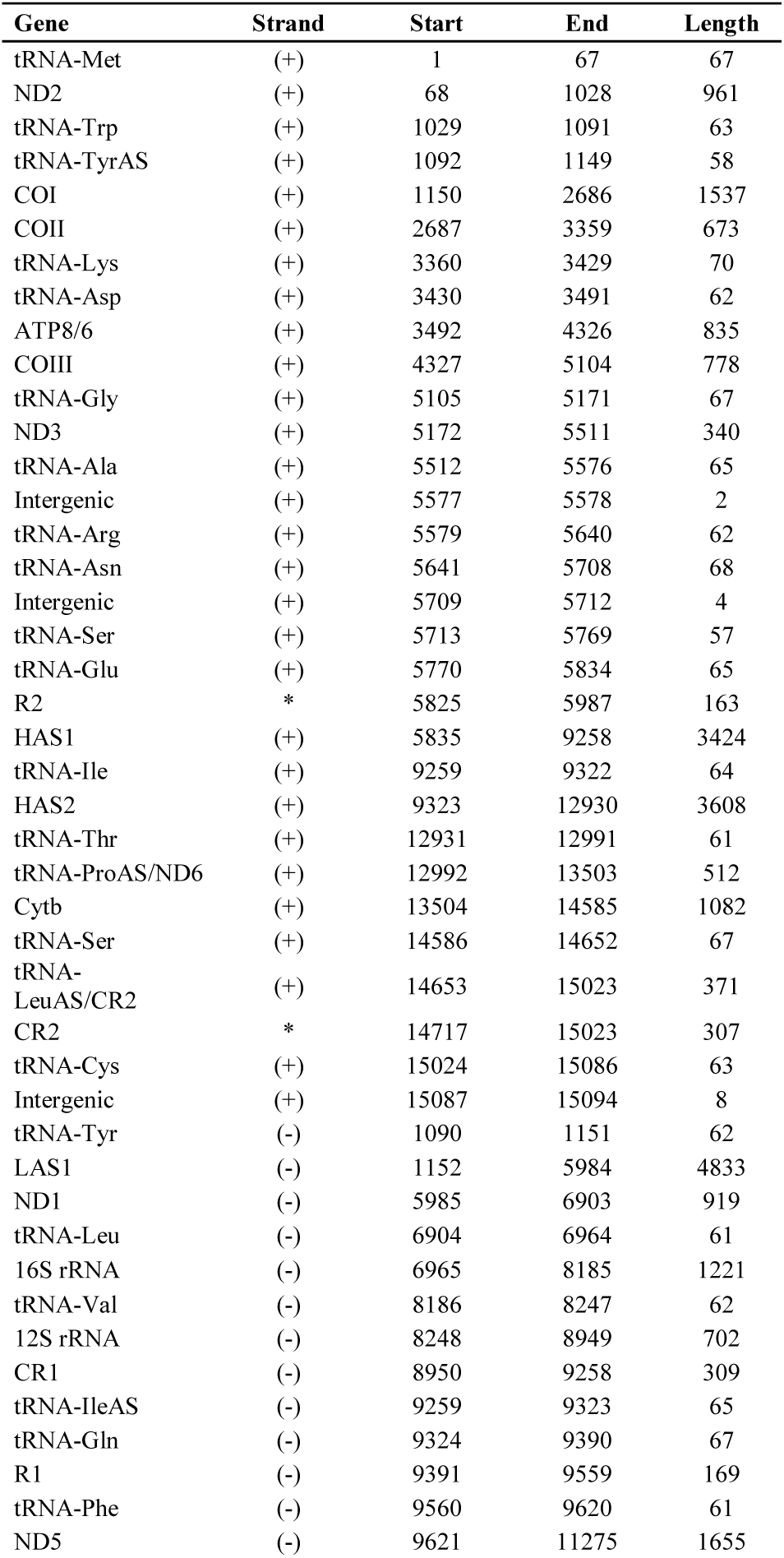
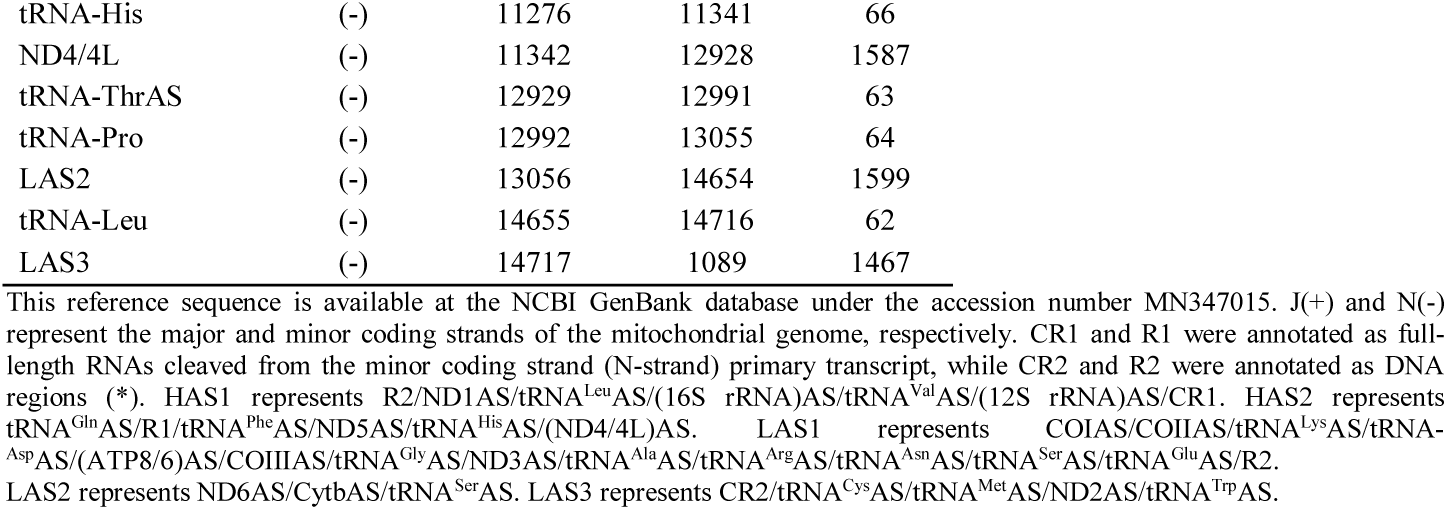
Precise annotations of the *D. silvarum* mitochondrial genome

Using precise annotations, we obtained two new findings in the *D. silvarum* mitochondrial tRNAs. The first involved six mitochondrial tRNA genes from which atypical tRNAs with no D-arm or an unstable T-arm were inferred [8]. One of tRNA^Ser^s and tRNA^Cys^ had no D-arms, while tRNA^Ala^, tRNA^Glu^, tRNA^Tyr^, and tRNA^Phe^ had unstable T-arms (**Figure 2C**). Another new finding was intergenic sequences between mitochondrial tRNA genes; mammalian mitochondrial genomes do not contain gaps, except for a novel 31-nt ncRNA [4]. Although these intergenic sequences were cleaved between their neighbouring tRNAs to form small RNAs (shorter than 10 bp), they were not likely to have biological functions. One typical example of an sRNA was A[U]_7_, between tRNA^Cys^ and tRNA^Met^ (**Figure 2C**). Based on these two findings, we found that 1×n STRs involved both intergenic sequences (e.g., A[U] _7_) and atypical mitochondrial tRNAs (e.g., [A]_5_ in tRNA^Cys^). Comparison of tRNA^Ser^ and tRNA^Cys^ suggested that tRNA^Cys^ with no D-arm had an [A]_5_ insertion that formed a large loop (**Figure 2C**). Given that the tRNA^Cys^ DNA sequence had too little evolutionary conservation to allow for an STR insertion, it proved a long-standing hypothesis that atypical tRNAs do not have biological functions.

R1 and R2 (**Figure 3**) were predicted to be two non-coding and non-transcriptional regions in the previous study [7]. However, they were proven to be transcribed on two strands in this study. The repeat units in R1 were reverse complements to those in R2 (**Figure 3B**). Our DNA-seq data showed that the copy numbers of R1 and R2 exhibited great diversity within an individual, which confirmed the finding from our previous study of the *E. fullo* mitochondrial genome [5]. As repeat units in R1 and R2 were reverse complements, we used PCR amplification (**Table 1**) combined with Sanger sequencing to further investigate R1 sequences in more than 100 individual ticks from five species (*D. silvarum, D. nuttalli, D. marginatus, D. niveus*, and *D. ushakovae*) and obtained the following results: (1) for each individual tick, the R1 sequence obtained using Sanger sequencing is actually a consensus sequence of a large number of heterogeneous sequences; (2) copy numbers were distributed between 2 and 5 for all studied repeat units, with one partial repeat unit counted as 1; (3) in total, three types of repeat units of R1 with lengths of 28, 34, and 44 bp (types 1, 2. and 3, respectively) were identified (**Figure 3B**) and noted as R_28_, R_34_ and R_44_; (4) in general, R1 sequences from ticks of one species were composed of repeat units of one type and R1 sequences from ticks of the same species from different places had different copy numbers; and (5) among all studied ticks, *D. nuttalli, D. niveus*, and *D. ushakovae* ticks only had R1 which was composed of the type 3 unit in their mitochondrial genomes, while *D. silvarum* ticks had R1 which was composed of the type 2 or 3 units. As for the *D. marginatus* ticks, most had R1 composed of the type 3 unit; however, a few had R1 composed of the types 1 and 2 hybrid units, noted as [R_34_]_l_-[R_28_]_m_-[R_34_]_n_, where l, m and n represent the copy numbers. The discovery of hybrid units suggested that mitochondrial DNA recombination can occur within an individual tick, resulting in the insertion of [R_28_]_m_ into [R_34_]_l+n_. This confirmed the identification of DNA recombination events in our previous study of the *E. fullo* mitochondrial genome [5]. In that previous study, the insertion of segments A and B into STR [R_87_] _l+m+n_ resulted in [R_87_]_l_-A-[R_87_]_m_-B-[R_87_]_n_.

### Discovery of a transposon-like element

A previous study identified a repeat unit dubbed the “tick box”—a degenerate 17-bp sequence motif directing the 3′ formation of ND1 and tRNA^Glu^ transcripts in all major tick lineages [7]. A large translocated segment (LT1) was reported spanning from ND1 to tRNA^Gln^ and the presence of the “tick box” motif at both ends of LT1 suggested its involvement in recombination events that are responsible for Metastriata genome rearrangements [7] (**Figure 1A**). Using precise annotations, LT1 was corrected to span R2, ND1, tRNA^Leu^, 16S rRNA, tRNA^Val^, 12S rRNA, CR1, tRNA^Ile^, tRNA^Gln^ and R1 (**Figure 3A**) in the reference genome. Given that nearly 50% of the human genome is contained in various types of transposable elements that contain repetitive DNA sequences [9], we hypothesized that LT1 is a transposon, with R1 and R2 as invert repeats (IRs) and genes from ND1 to tRNA^Gln^ as insert sequences (ISs). To validate our hypothesis, we detected structural variation (**Materials and Methods**) in the *D. silvarum* mitochondrial genome to determine the occurrence of LT1 translocation events. The results proved the occurrence of LT1 inversions within an individual tick (**Figure 3A**).

As the occurrence of LT1 inversions is rare, 4.1 Gbp DNA-seq data were generated to cover ∼427,247 × (4 .09 Gbp/9.58 Kbp) of L1 in the *D. silvarum* mitochondrial genome to detect the LT1 inversions. As the dominant copy number is five for both R1 and R2, we used 34*5 STR to represent R1 and R2 in the *D. silvarum* mitochondrial genome. Thus, R1 and R2 in *D. silvarum* are ∼170 bps long (34*5 STRs), which is longer than the reads in the 2 × 150 bp DNA-seq data. We had to sequence the same library using 2 × 250 bp sequencing to validate the reference genome and the LT1 inversion (**Materials and Methods**). The substantial diversity in R1 and R2 copy numbers within an individual tick rendered great diversity in LT1. However, we did not obtain full-length sequences of LT1 due to sequence-length limitations in the DNA-seq data. Therefore, we were unable to determine whether R1 and R2 had the same copy numbers within one LT1.

### STR variation in the mitochondrial genomes

By mapping DNA-seq data to the *D. silvarum* mitochondrial genome, variation detection was performed to report two types of DNA variation (**Materials and Methods**)—SNPs and small insertions/deletions (InDels). Out of our expectation, all of the detected variants exhibited changes in the copy numbers of STRs caused by InDels of one or more entire repeat units, while SNPs were not detected within a *D. silvarum* tick. We deemed these copy number changes as STR variation and defined the STR position as the genomic position of the first nucleotide of the reference STR. For example, [G]_8_ was designated as the reference STR at position 1810, because it occurred most frequently in mtDNAs within one individual tick (**Table 3**); the alternative alleles of [G]_8_ included [G]_6_, [G]_7_, [G]_9_, and [G]_10_. Importantly, it was found that almost all of the STRs, particularly those with copy numbers greater than 5, had variants. The detection of STR variation was reliable, based on the following reasons: (1) the PCR amplification and high-depth DNA sequencing produced a high signal-to-noise ratio in the detection of DNA variation; (2) the Illumina sequencer does not generate InDel errors in the DNA-seq data; (3) the alternative allele ratios (**Materials and Methods**) at 20 STR positions are significantly higher and the highest ratio reached was ∼33% at position 3441 (**Table 3**); (4) it was impossible for sequencing or alignment errors to result in 2-bp InDels in 2×n STR (e.g., [TA] _9_); (5) the same STR variation caused diversity within and between individuals, e.g., [TA]_9_ in our genome (**Table 3**) and [TA]_5_ in NC_026552.1; (6) STR variation was detected in other animal species, including *E. fullo* [5], mouse and human [6].

Using a very strict parameter (**Materials and Methods**), we selected 20 STR positions for further analysis (**Table 3**). These STRs resulted in highly diversity between *D. silvarum* individuals. They can be used as DNA markers in order to discriminate between individuals. Almost all of them were composed of A or T, except [G]_8_ at position 1810. STRs at 18 positions were 1×n STR and those at the other 2 positions were 2×n STR. All of the reference STRs and their variants had copy numbers greater than 5. Among all of the variants, 1-bp InDels occurred more frequently than longer ones. In particular, 13 STR positions resided in the protein-coding genes and the variants at these positions resulted in frameshift mutations in the proteins, which can produce deleterious effects [10]. For example, the COI gene has a 1524-bp Coding Sequence (CDS) for a 308-aa protein. The variant [G]_9_ at genomic position 1810 (**Table 3**) can result in a 765-bp CDS for a 255-aa truncated COI protein. Thirteen STR positions had identical sequences, exhibiting high evolutionary conservation; however, the other seven STR positions in the tRNA and rRNA genes exhibited variation between individuals. For example, [A]_10_, [TA]_9_, [T]_9_ and [T]_10_ at positions 5524, 7324, 8042 and 8243 in our mitochondrial genome changed to [A]_8_, [TA]_5_, [T]_8_ and [T]_6_ at positions 5524, 7324, 8042 and 8243 in NC_026552.1 (**Table 3**). This finding inspired us to investigate if animal cells have mechanisms to remove mitochondria containing deleterious mutations or inhibit the expression of the deleterious variants, as they can cause loss of function or diseases. However, we did not obtain high-depth RNA-seq data for *D. silvarum*. We had to compare the STR variants in the *E. fullo* mitochondrial genome [5] at the DNA level and RNA levels, using the DNA-seq and RNA-seq data (SRA: SRP174926). However, we found no significant differences between them. This suggested that deleterious STR mutations can irreversibly change the proteins made from their mRNAs and that mitochondria containing deleterious mutations can accumulate in cells.

**Table 3.**
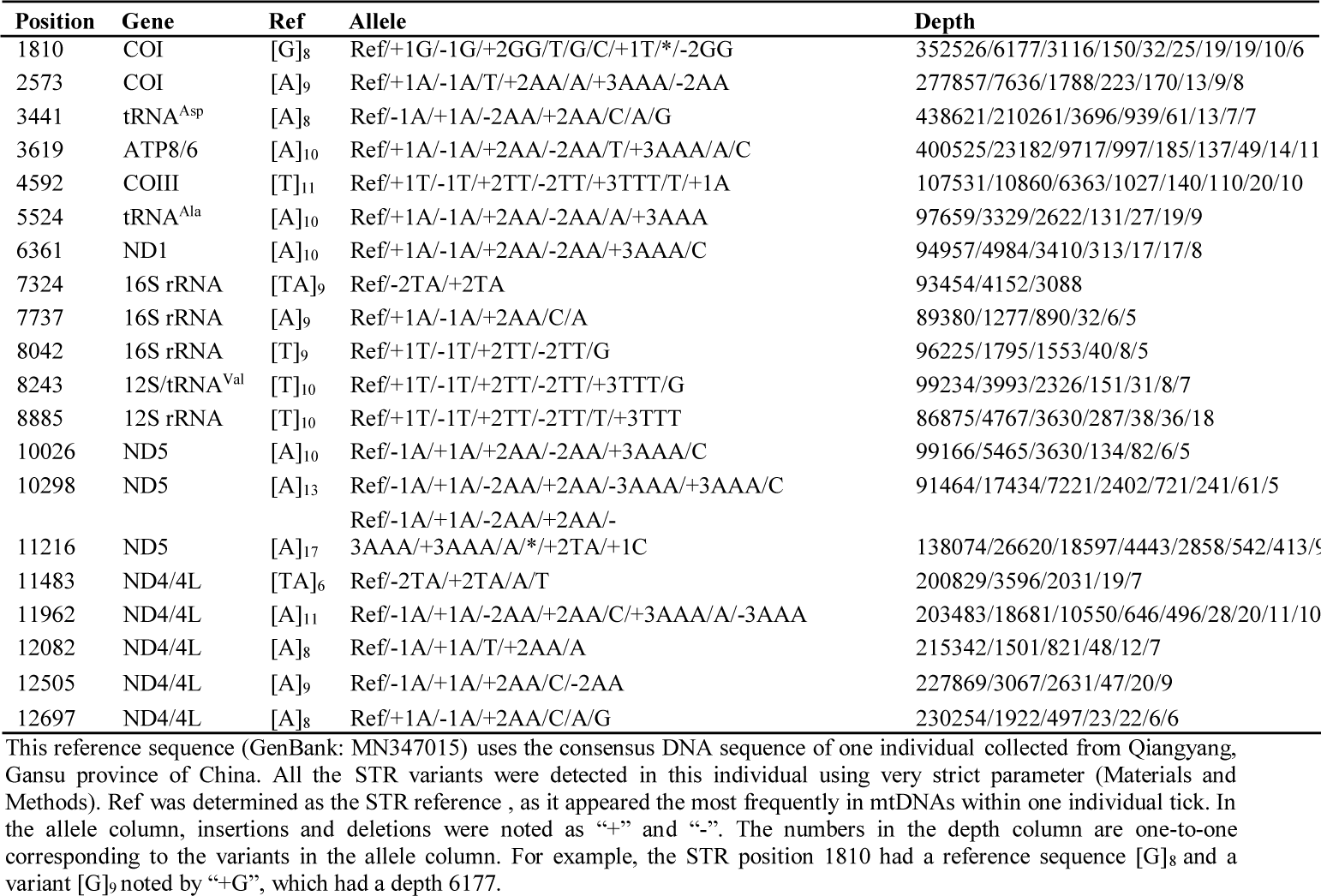
Mitochondrial STR variation within an *D. silvarum* individual

## Conclusion and Discussion

In this study, we used long-PCR amplification combined with NGS to obtain complete mitochondrial genomes of individual ticks and performed precise annotation of these genomes. Using these precise annotations, we achieved our research goals (**Introduction**) along with two new findings that merit future study. The discovery of a transposon-like element may shed light on the mechanisms of mitochondrial gene order rearrangement and genomic structural variation, especially with additional data from more tick species. The second finding paved the way to an eventual understanding of the mechanisms of mitochondrial DNA variation. The comparison between interindividual and intraindividual variation showed that STRs with shorter repeat units (e.g., 1×n STRs) and STRs with longer repeat units (e.g., 34×n STRs) had the same variation pattern. Our findings will encourage reconsideration of the importance of STRs as well as a unified study of STR variation with shorter and longer repeat units in both nuclear and mitochondrial genomes.

SNPs, accepted as the most common type of genetic variation, play a dominant role in studies in almost all biological fields; however, research of STRs has been limited to specialized fields. In particular, the use of SNPs is becoming dominant in the studies of mitochondria, e.g., mitochondrial heterogeneity. After further analysis of tick, insect, mouse and human DNA-seq and RNA-seq data, we found that a large number of STR variants had been missed in variation detection, as most research has focused only on the detection of SNPs using WGS data. Although SNP platforms are higher throughput and more cost-effective for genome scans, STRs remain highly informative measures of DNA variation for linkage and association studies. Using PCR combined with very high depth DNA-seq data, we were able to accommodate three gaps in genome alignment and still ensure detection accuracy. We also proved that WGS data can be used to detect STR variation with careful adjustment of parameters (**Materials and Methods**). Future studies are necessary on the origins, mutation rates, and effects of STR variation in, but not limited to, animal mitochondrial genomes.

Both 1×n STRs and 34×n STRs had the same variation pattern; 34×n STRs can be produced by DNA recombination, however, the cause of 1×n STRs remains unclear. One possible cause is replication slippage [11]. In this theory, DNA polymerase causes mismatches between DNA strands while being replicated during meiosis. It can then slip while moving along the template strand and continue on the wrong nucleotide. Another possible cause could be point mutations, based on a study comparing human and primate genomes [12]. Both replication slippage and point mutations can not be used to explain the occurrence of m×n STRs (m=2, 3, 4, 5 …). Using the *E. fullo* full-length transcriptome data [5], we also detected multiple STR variations. The PacBio sequencer can be used as an *in vitro* system to test the replication slippage theory. However, DNA polymerase in the PacBio sequencer and the signal detection system can also cause InDels. A comparison between Illumina and PacBio data can be performed to determine whether DNA polymerase in the PacBio sequencer causes STR variation.

Unique DNA sequences in a genome exhibit a very low variation/mutation rate (approximately 10^−9^ nt per generation), whereas the mutation rates in STRs are several orders of magnitude higher, ranging from 10^−6^ to 10^−2^ nt per generation for each locus [13]. Direct estimates of STR mutation rates have been made in numerous organisms from insects to humans, e.g., *Schistocerca gregaria* 2.1 × 10 ^−4^ [14]. In this study, we only detected STR variation within an individual animal and found that the alternative allele ratio was distributed from less than 0.01% to ∼33 %. This suggested that STR variation occurrences varied significantly along the genomes such that mutations concentrated at certain positions (e.g., 20 STR positions for the *D. silvarum* mitochondrial genome). Future studies must be performed to estimate STR mutation rates of mitochondria to test if they have correlations with life expectancy of different animals, using individuals at different developmental stages.

It is well accepted that many STRs are located in non-coding DNA and are biologically silent, while others are located in regulatory or even coding DNA. STRs located in regulatory, intron and transposon regions are beyond the scope of this study. However, our studies showed there was no significant differences between the alternative allele ratios of STR positions in the protein-coding gene regions and those in tRNA and rRNA gene regions in tick mitochondrial genomes. Particularly, mitochondria containing deleterious mutations can accumulate in cells and deleterious STR mutations irreversibly change the proteins made from their mRNAs. This suggested that deleterious STR mutations in mitochondria cause aging and diseases, known as Tandem Repeat Disorders (TRDs). Huntington’s disease, as one of the famous TRDs, occurs in the context of expanded glutamine [CAG]_n_ repeats. Several other human diseases have also been linked to mitochondrial STR variation. Another famous example—breast cancer (BC)—has been linked to D_310_ variation [15]. The telomeres at the ends of the chromosomes consist of [TTAGGG]_n_ in vertebrates. Thus, both telomeres and mitochondria are linked to ageing/senescence by STR variation.

## Materials and Methods

Individual ticks from five species (*D. silvarum, D. nuttalli, D. marginatus, D. niveus* and *D. ushakovae*) of the genus Dermacentor were collected from different places in china and were identified using a stereoscopic microscope according to [16]. Total DNA was isolated from individual ticks using QIAamp DNA Mini Kit (Qiagen, Germany), following its protocol. Long-PCR amplification of each entire mitochondrial genome in L1, L2, L3 and L4 was performed (**Figure 1A**) using Long PCR Mix (Sino-Novel, China) and the PCR reaction mixture was incubated at 95°C for 3 min, followed by 34 PCR cycles (30 s at 98°C, 30s at 55°C and 5 min at 68°C for each cycle) with the final 5 min extension at 72°C, while PCR amplification of short segments (CR1, CR2, R1 and R2) was performed using Taq PCR Mix (Sino-Novel, China) and the PCR reaction mixture was incubated at 95°C for 3 min, followed by 34 PCR cycles (30 s at 95°C, 30s at 55°C and 1 min at 72°C for each cycle) with the final 5 min extension at 72°C. All the primers and PCR reaction conditions are listed in **Table 1**. The amplified L1, L2, L3 and L4 of *D. silvarum* and L3 and L4 of *D. nuttalli* and *D. marginatus* were used to construct ∼250-bp-size libraries, respectively and sequenced using 2*150 bp paired-end strategy on the Illumina HiSeq X Ten platform. The amplified L1 of *D. silvarum* was also used to construct one ∼350-bp-size library and sequenced using 2*250 bp paired-end strategy on the Illumina HiSeq 2500 platform (**Figure 1B**). On average, 2 Gbp DNA-seq data was obtained for L1, L2, L3 and L4, respectively, thus, 4 Gbp data was obtained for each mitochondrial genome. In total, 10 runs of DNA-seq data (L1 of 2*250, L1, L2, L3 and L4) of *D. silvarum* were submitted to the NCBI SRA database under the project accession number SRP178347. The assembly of tick mitochondrial genomes was performed using the software MIRA 4.0.2. The complete mitochondrial genome sequence of *D. silvarum* is available at the NCBI GenBank database under the accession number MN347015. However, precise annotations of the *D. silvarum* mitochondrial genome need be seen in **Table 2**, as the NCBI GenBank database is not able to accept this new annotation format. The amplified short segments were sequenced using Sanger sequencing (**Table 1**). Using the software BWA, DNA-seq reads were aligned to the *D. silvarum* mitochondrial genome with parameters (-n 3 -o 1 -e 2 -l 15 -k 1) to detect DNA variation. The STR variants (**Table 3**) were detected by single-end alignment using the 2*150 bp DNA-seq data of L1 and L2, and then, validated using the 2*150 bp DNA-seq data of L3 and L4. The alternative allele ratio was calculated by the depth of allele divided by the total depth on the genomic position. The 20 STR positions were selected using a strict parameter, which required the alternative allele ratio above 1%.

Total RNA was isolated from ticks to construct sRNA-seq libraries, following the protocol [17] and these libraries were sequenced using 50-bp single-end strategy on the Illumina HiSeq 2500 platform. In total, two, two and two runs of sRNA-seq data from *D. silvarum, D. nuttalli* and *D. marginatus* were submitted to the NCBI SRA database under the project accession number SRP178347. Total RNA was isolated from *D. silvarum* to construct one PacBio full-length cDNA library, following the protocol [2] and this library was sequenced on the PacBio Sequel sequencer to obtain the PacBio full-length transcriptome data. The cleaning and quality control of sRNA-seq and the PacBio full-length transcriptome data were performed using the pipeline Fastq_clean [18]. Using the software bowtie, sRNA-seq reads were aligned to the *D. silvarum* mitochondrial genome with one mismatch for precise annotation. To confirm the precise annotation of the *D. silvarum* mitochondrial genome, the PacBio full-length transcriptome data was used to follow the same procedure as [5]. The structural variation detection was performed following the protocol as [19]. Statistics and plotting were conducted using the software R v2.15.3 the Bioconductor packages [20].

## Acknowledgments

We appreciate the help equally from the people listed below. They are Professor Defu Chen, Guoqing Liu, Dawei Huang, Yanqiang Liu, Associate Professor Bingjun He, Qiang Zhao and the graduate student Xiufeng Jin, Haishuo Ji, Guangcai Liang from College of Life Sciences, Nankai University. We also appreciate the help equally from Professor Jishou Ruan from School of Mathematical Sciences, Nankai University. We would like to thank Ellen_7 from Editage (www.editage.cn) for English language editing.

